# Semi-Automatic Segmentation of *Pseudomonas koreensis* and *Escherichia coli* for Bacterial Growth Characterization

**DOI:** 10.1101/2024.08.15.608040

**Authors:** Diana A. Alvarado-Ruiz, Keny Ordaz-Hernández, Lourdes Díaz-Jiménez, Grecia L. Lara-Cadena, Roberto González-López, Gregorio Vargas-Gutiérrez, Mario Castelán

## Abstract

Bacterial characterization is a crucial discipline within microbiology. Given the manual and labor-intensive nature of this task, our aim is to introduce a semi-automatic segmentation method that enhances efficiency while preserving the rich details of bacterial colonies. We propose using the k-means clusterization algorithm to analyze and segment images of bacterial cultures, specifically those of *Pseudomonas koreensis* and *Escherichia coli*. Unlike existing methods that focus primarily on colony counting, our approach emphasizes morphological characterization. In some bacterial cultures, colonies are not well-defined, making manual counting or other automated counting methods unfeasible; i.e. the bacterial growth area is not easily identifiable, thus precise growth tracking is not feasible. Our method enables bacterial growth characterization even in these cases. Our computer vision system identifies and quantifies the diverse morphologies within *P. koreensis* and *E. coli* cultures, determining their relative occupancy in an image. Our approach provides valuable insights into the composition, growth patterns, and developmental stages of bacterial colonies, designed to assist both novice and expert microbiologists in bacterial analysis.

## Introduction

Bacterial characterization is an important activity with significant implications across various sectors, including industrial and pharmaceutical applications. Characterizing bacterial growth requires methodologies that offer rapid results without sacrificing accuracy or the well-being of analysts. Microbiological characterization methods, as detailed by Ramirez [1], range from traditional techniques like colony counting [2], which provides a general overview of microbial communities [3], [4], and [5], to more advanced technologies designed to make these processes more flexible [6], [7], [8], [9], [10] and [11]. Despite technological advancements [12], manual counting of bacterial colonies remains prevalent among some laboratory workers. This method is not only tedious [13] and time-consuming [14], but it also increases the risk of errors and health issues such as visual stress, which can lead to fatigue, frustration, and ocular problems [15]. In addition to this, the traditional metric of Colony-Forming Units (CFU) is often inadequate for characterizing growth patterns of colonies with complex or irregular morphologies. For these diverse and non-standard shapes, a more morphologically flexible unit of measurement is essential to thoroughly understand the sequential development and growth of colonies.

To address these challenges, we propose a semi-automatic procedure for bacterial growth characterization that utilizes computer vision via the k-means algorithm for analyzing and segmenting images of bacterial cultures, specifically *P. Koreensis* [16] and *E. Coli* [17]. This innovative method significantly reduces the required time, compared to manual counting, and shifts the focus from traditional CFU counting to characterizing the percentage of occupancy within the culture. By doing so, our strategy enables the detection and analysis of various morphologies present, providing critical insights into the growth, interaction, and developmental behavior of the colonies over time.

## Materials and methods

This section presents the principal phases of the proposed procedure, as depicted in Figure 1. The process begins with the cultivation of bacterial samples, progressing through to the application of the semi-automatic colony characterization using the k-means algorithm [18], [19]. Key steps include the preparation of the bacterial cultures, the selection of representative samples for analysis, and the setup of the necessary software and hardware installations. Additionally, this stage involves creating and curating a database that integrates the segmented and analyzed data from the images.

**Figure 1.**
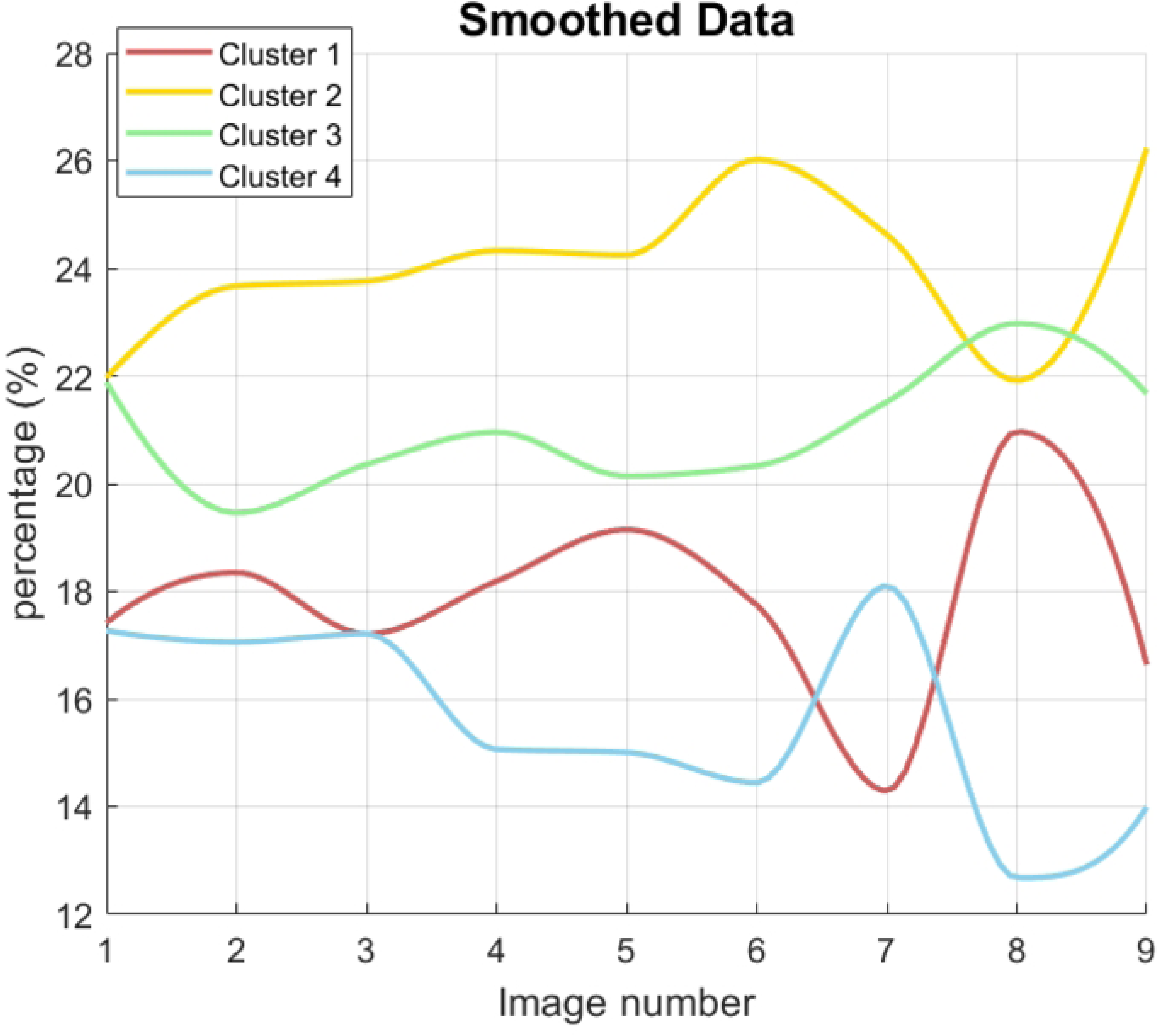
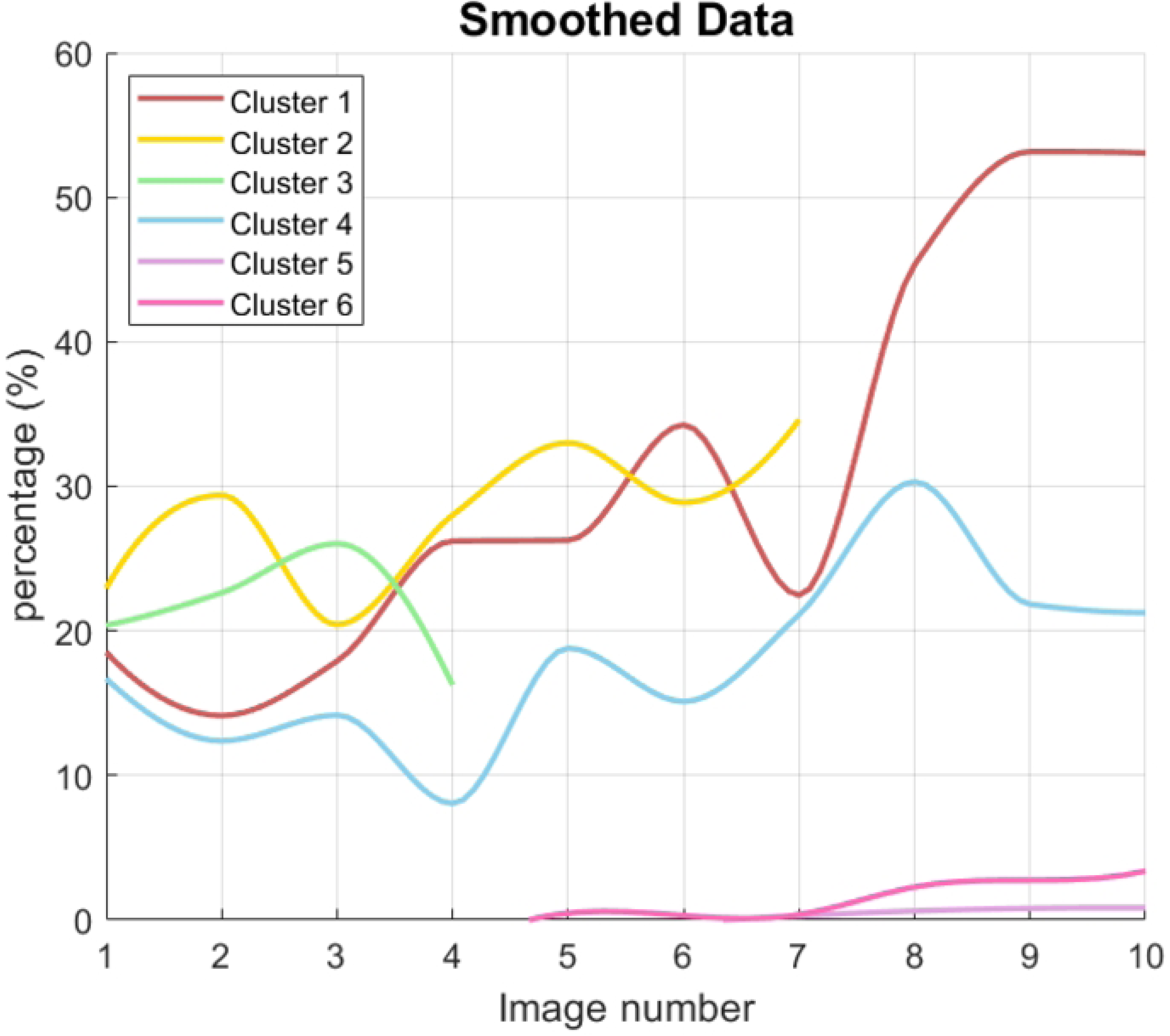
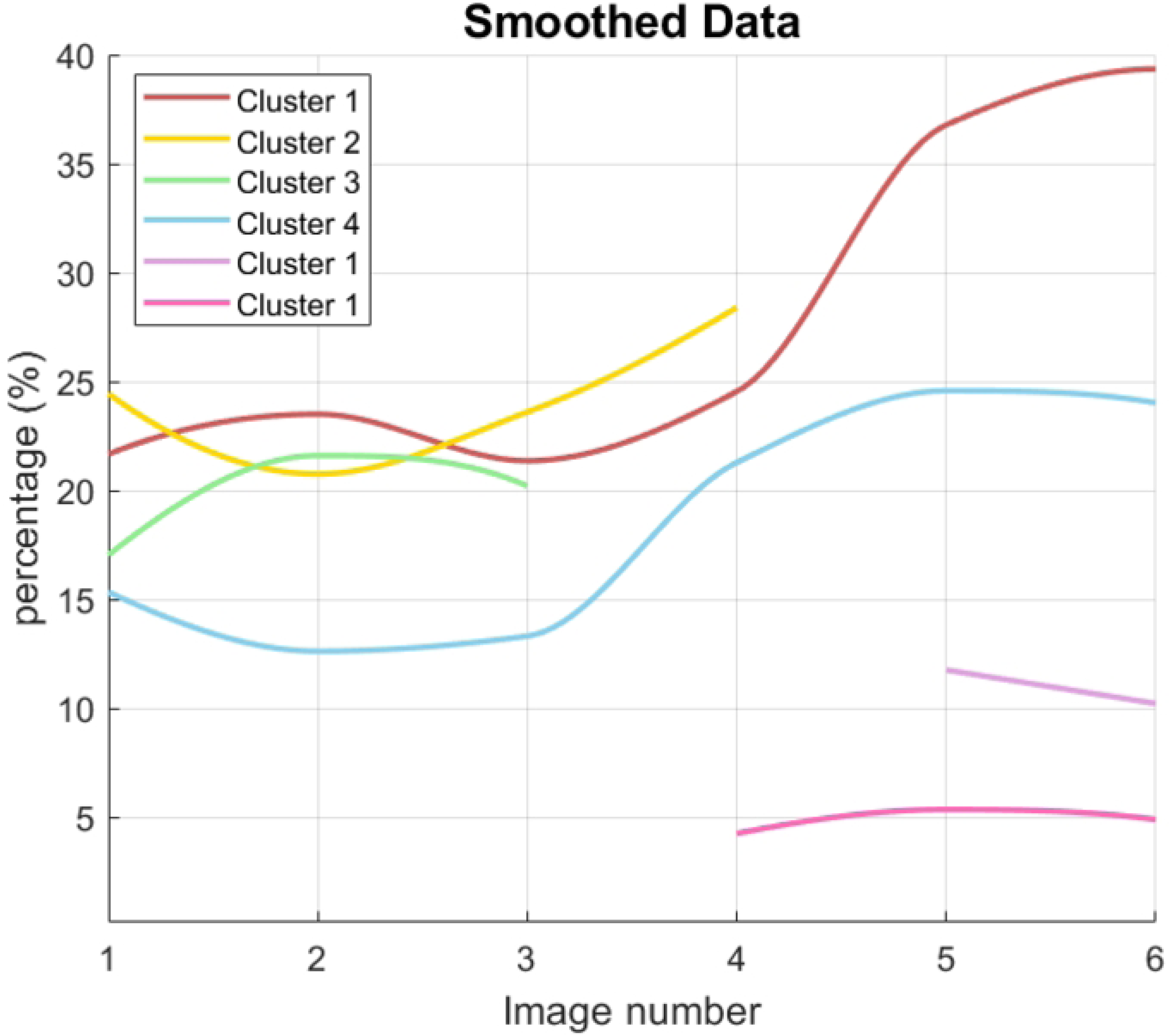
Main stages of the proposed procedure. The figure depicts six steps, from bacterial culture preparation through bacterial occupancy determination.

### Bacterial culture

The bacterial species utilized in this study were *P. koreensis* and *E. coli*. The culture process for *P. koreensis* involved the use of 60*×*15mm Petri dishes filled with cetrimide agar. This procedure adhered to the established protocols as detailed in [20] and [21]. In contrast, the cultivation of *E. Coli* was conducted in larger 90*×*15mm Petri dishes using nutrient agar, following the guidelines provided by [22].

### Image acquisition and database

For image acquisition, a specialized setup was constructed as shown in Figure 2. This system comprises a Pi Camera Rev. 1.3, which is mounted on a wooden arm positioned above the colony counter device. The camera is connected to a Raspberry Pi 3 model B board, which facilitates image capture and processing. The Raspberry Pi board is further linked to a keyboard (serving as the capture device) and a screen (used as the display device). The images captured using this setup have a resolution of 2592*×*1944 pixels. To facilitate the image-saving process, each image was automatically saved under a generic name.

**Figure 2.**
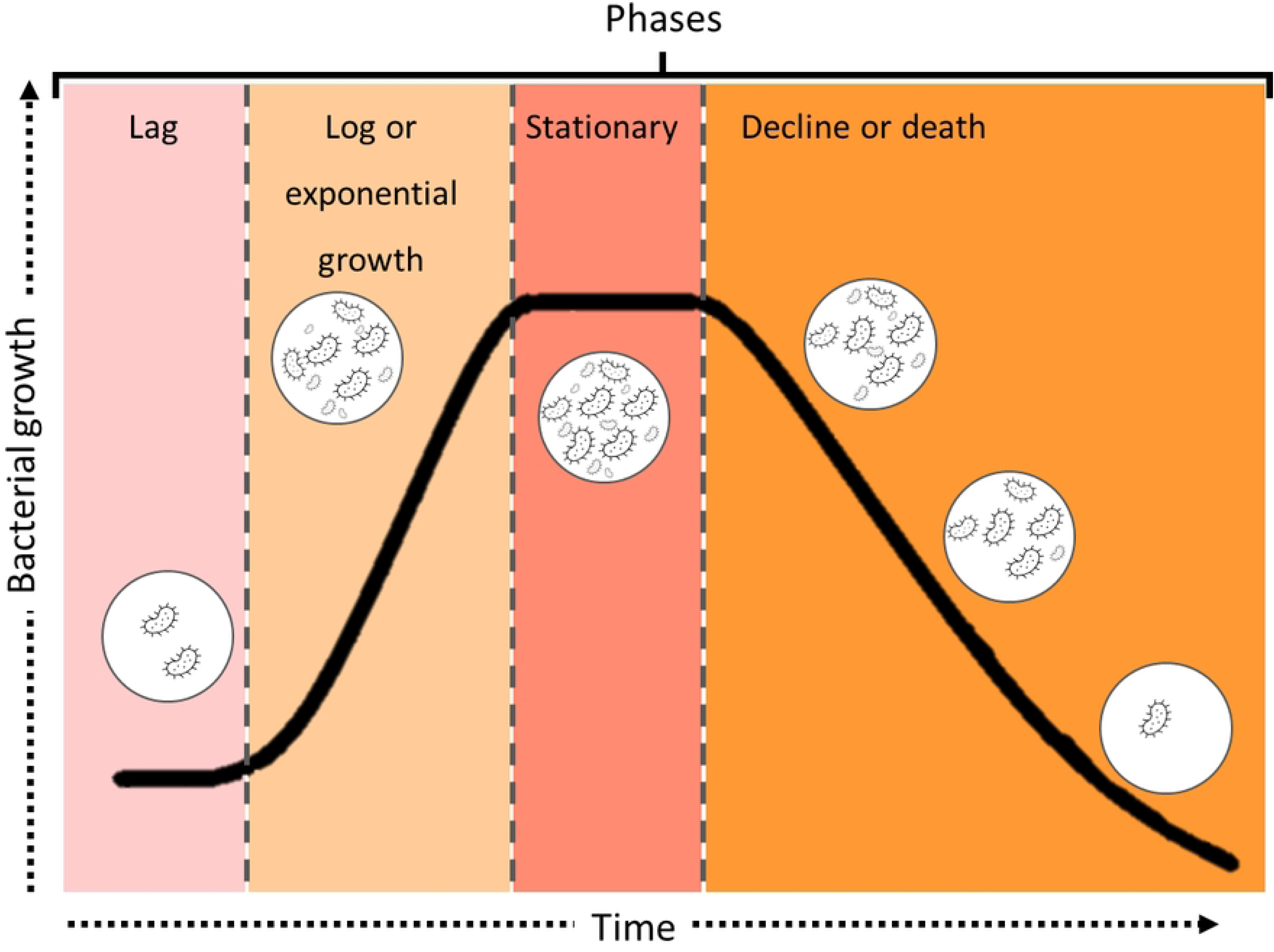
Image acquisition platform. Installed in the Waste Revaluation Laboratory of the Sustainability of Natural Resources and Energy Postgraduate Program at CINVESTAV-Saltillo

To minimize sample transportation and manipulation time, the imaging platform was installed on a table situated above the incubator where the cultures were stored, ensuring proximity and easy access during the experimental process. To achieve optimal and consistent lighting conditions, several precautions were implemented. All external lights were turned off and window blinds were closed to eliminate reflections on the colony counter’s magnifying glass and the Petri dishes. This setup helped maintain the integrity of the image captures by preventing any light interference that could affect the accuracy of the results. The timing between sample captures was meticulously scheduled over three consecutive days, with each sample undergoing five image captures. From this process, three distinct cases were compiled, yielding a total of ten, six, and nine images respectively, after data cleaning was conducted. The first two cases involved *P. koreensis* cultures, while the third case pertained to an *E. Coli* culture, providing a diverse dataset for analysis. In Table 1, the specifications include details of each case, bacteria type, number of images, and the daily schedules.

**Table 1.** Case Descriptions and Daily Schedules.

| Case | Bact. Type | Schedule |  |  | Images (total) |
| --- | --- | --- | --- | --- | --- |
|  |  | Day 1 | Day 2 | Day 3 |  |
| 1 | *P. Koreensis* | 14:30h | 12:20h | 13:30h | 10 |
|  |  | 16:30h | 14:40h | 15:30h |  |
|  |  | 18:30h | 16:30h | 17:30h |  |
|  |  |  | 18:30h |  |  |
| 2 | *P. Koreensis* | 17:15h | 10:00h | 12:00h | 6 |
|  |  | 19:30h | 17:15h | 17:30h |  |
| 3 | *E. Coli* | 09:00h | 09:00h | 09:00h | 9 |
|  |  | 11:00h | 11:00h | 11:00h |  |
|  |  | 17:00h | 17:00h | 17:00h |  |

Table notes: Details include case-specific information, bacteria type, daily scheduling details, and total images.

### Image processing and k-means characterization

This phase of the project was carried out using Python version 3.9.12, incorporating essential libraries such as OpenCV for image processing, NumPy for numerical operations, and Matplotlib for data visualization, ensuring a robust environment for handling the data processing tasks.

The initial step in the bacterial characterization process involves image pre-processing, which is critical for accurate analysis. The process begins with the detection of the Petri dish in the image using the Hough Transform algorithm, a feature of the OpenCV library. This algorithm identifies circular objects, effectively isolating the Petri dish. Once detected, the first Region of Interest (ROI) [23] is established around the Petri dish. A binary mask [24] is then applied to the original image to narrow down the working area, focusing exclusively on regions where bacterial growth occurs.

Following this, a second ROI is created. This step involves the application of the k-means clustering algorithm to categorize the image into three distinct clusters [25], [26], [27]: the background, the Petri dish, and the agar/bacterial culture. This segmentation effectively highlights and separates each component, simplifying further analysis. A second binary mask is applied based on the first ROI, further refining the area under study.

The final segmentation involves re-applying the k-means algorithm with an adjusted number of clusters based on the elbow method [28] and visual feedback. This technique helps determine the optimal number of clusters by evaluating the trade-off between the number of clusters and the within-cluster sum of squares, allowing for a detailed yet efficient segmentation. This step is crucial as it ensures that the segmentation captures all relevant bacterial morphologies without omitting critical details, thereby facilitating a comprehensive understanding of the bacterial culture in each image.

Following segmentation, the images are recolored to enhance the visibility of distinct clusters within the Petri dishes. This enhanced visualization is demonstrated in Fig. 3, which displays Case 1. In this figure, ten images of *P. koreensis* are shown in their original color (top row) and k-means recolored (bottom row). It is noteworthy that the original images begin to show colony formation suitable for counting only by the third day. In contrast, the segmented and recolored counterparts facilitate the differentiation of various bacterial morphologies coexisting within the same sample, even in the early stages, aiding the experts in conducting a more detailed analysis.

**Figure 3.**
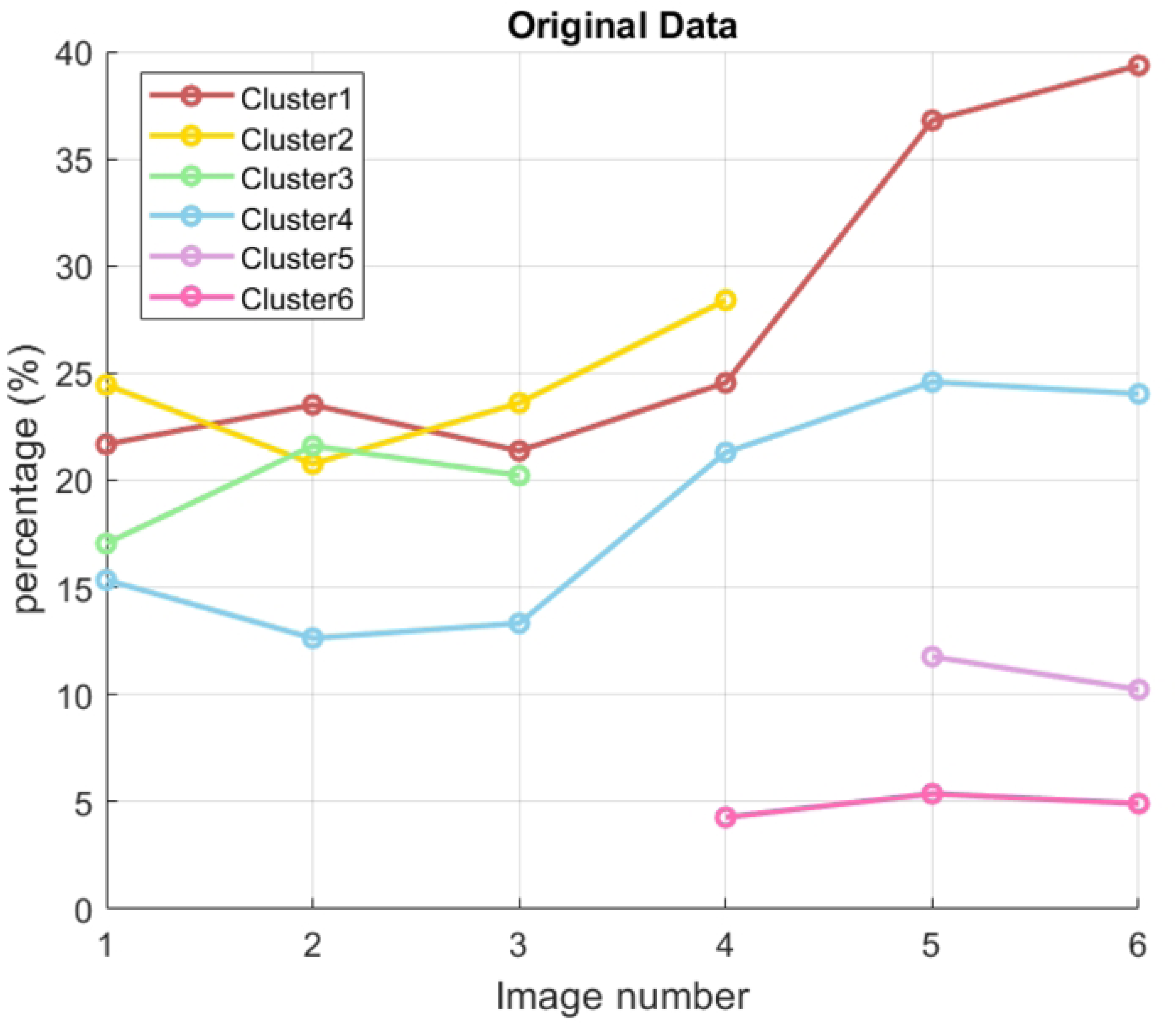
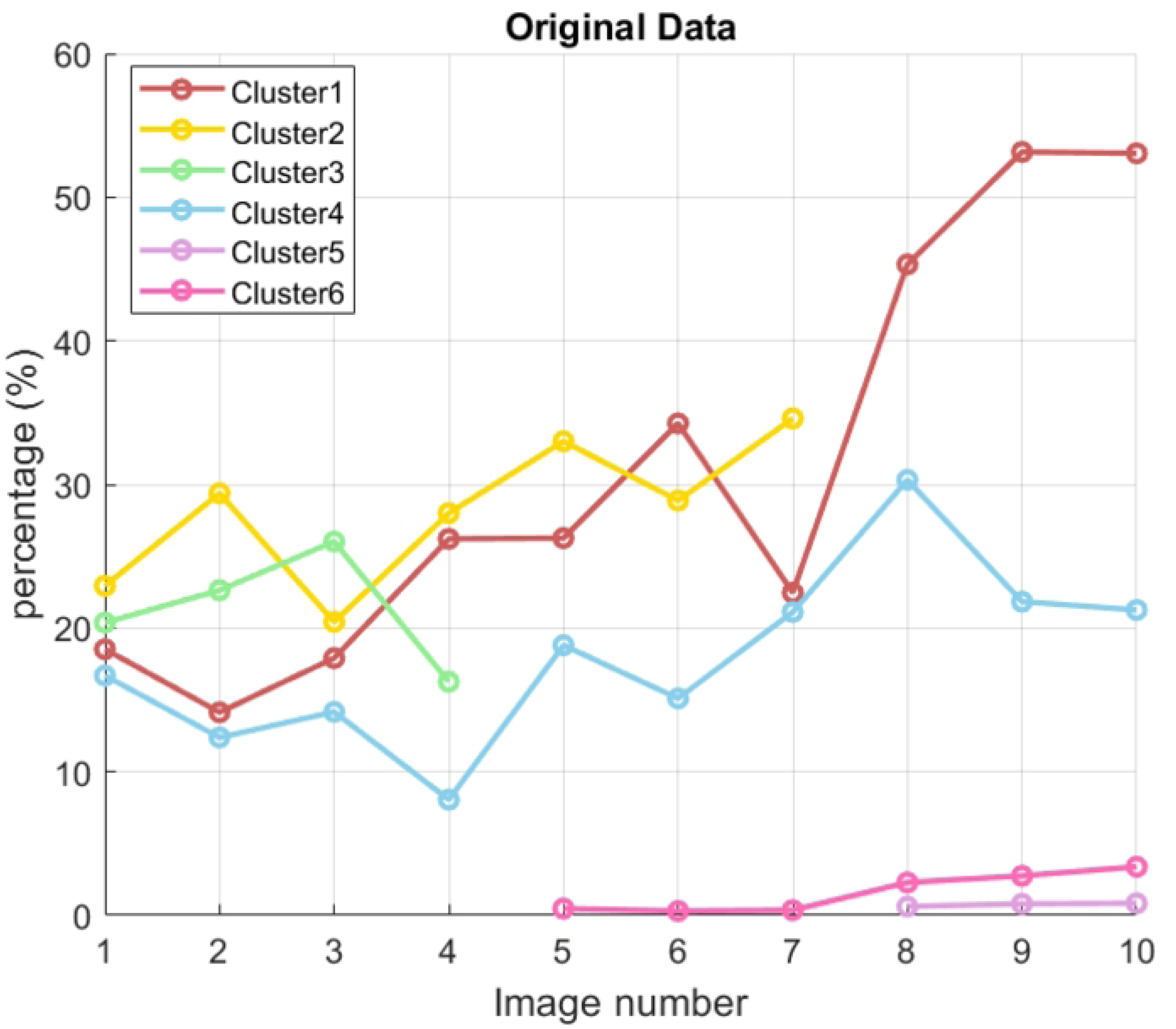
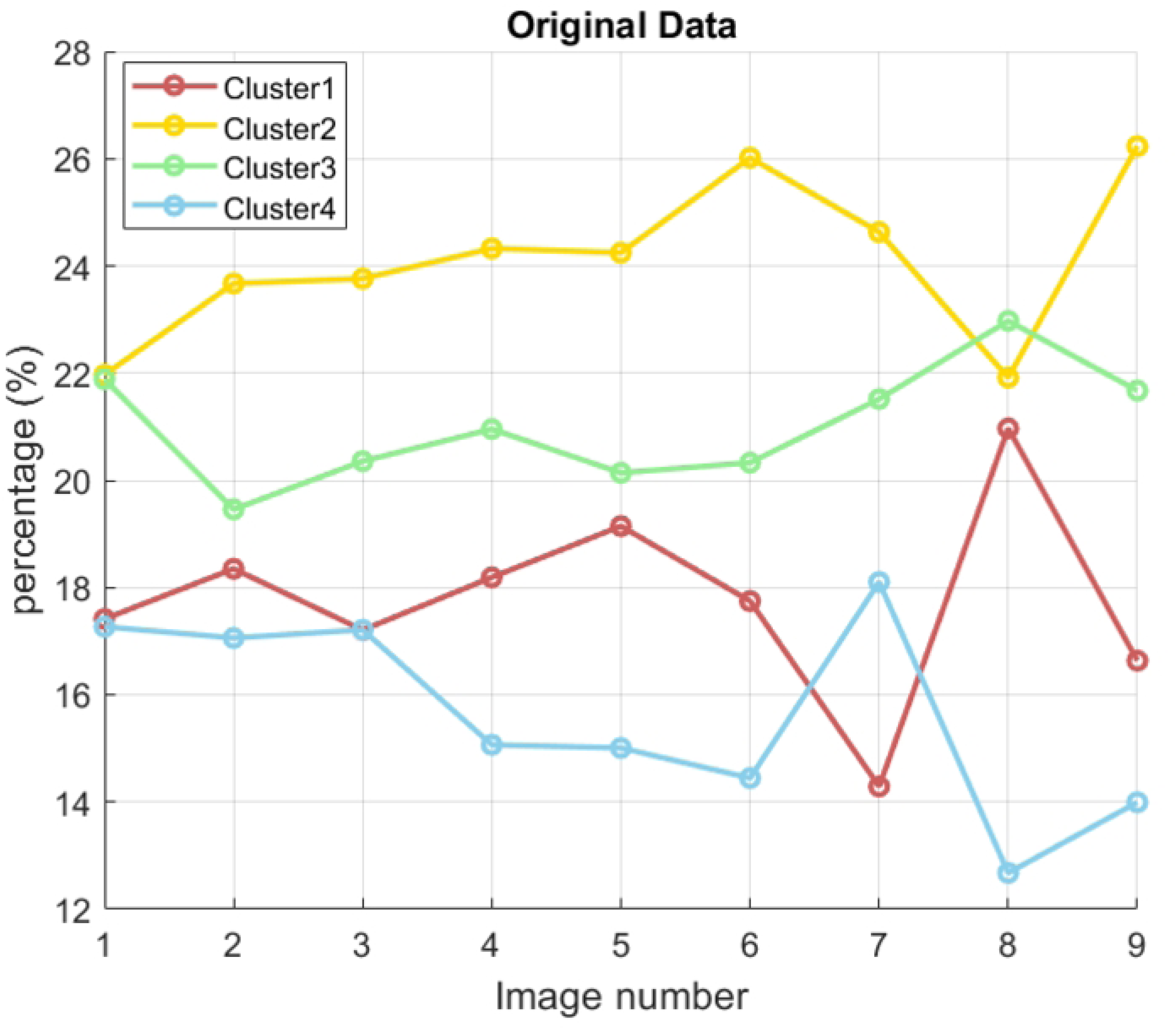
Original and recolored images. Ten images of *P. koreensis* (Case 1) are shown before (first two rows) and after having being k-means recolored (last two rows).

To further refine the analysis, relationships between the clusters are established automatically based on the Euclidean distance between their centroids. This metric helps determine which centroids—and consequently, which clusters—are closest to each other, providing a clearer understanding of the spatial relationships of the different formations within the Petri dish. Figure 4 visually compares these automatic relationships for the three cases in the experiment. Notably, the automatic calculation of clusters allows for an even more detailed visual examination of bacterial growth compared to the previous recolorization stage.

**Figure 4.**
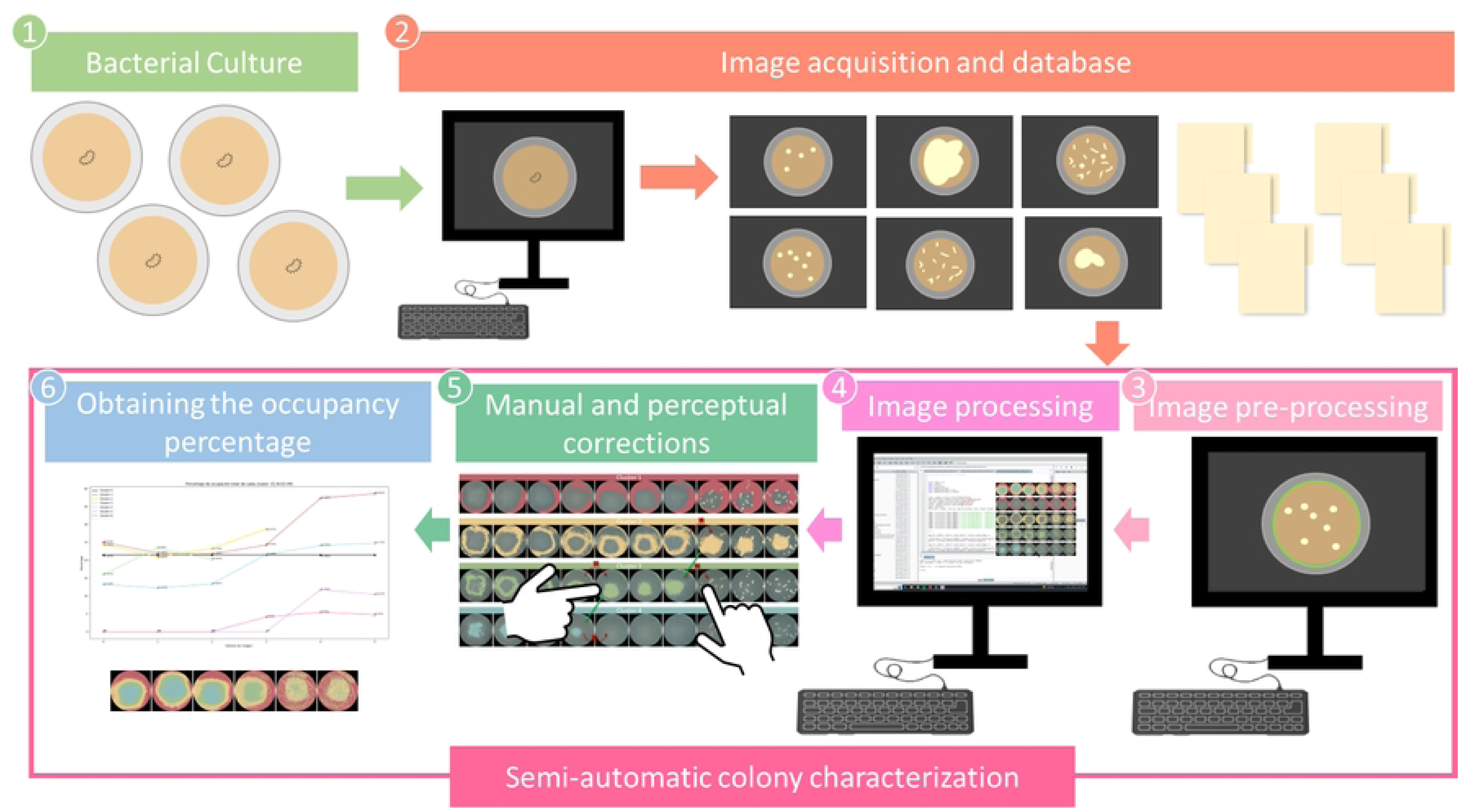
Recolored clusters for each case. Three cases are presented: Case 1 and 2 for *P. Koreensis* and Case 3 for *E. coli*. Recolorization results allow the expert for a visual analysis of the cluster that may lead to further manual corrections.

Once the clusters are obtained, it becomes easier to observe how different clusters evolve over time. For each of the three cases (*P. koreensis* in the first two panels and *E. coli* in the last panel), each row in the panel represents different clusters, while each column represents variations over time. The temporal variations remain consistent across columns within each panel.

### Manual and perceptual corrections

Although the previous stage involves automatic segmentation and cluster correspondence, a further stage is needed where the expert selects clusters that are not well corresponded. This is what we refer to as semi-automatic analysis, involving human participation to determine the final number of clusters in case new ones emerge. The semi-automatic nature of the proposed procedure arises from the fact that the results obtained through automatic clustering may not always precisely align with the morphological characteristics detected in the samples. Therefore, expert intervention is essential to verify and adjust the relationships suggested by the algorithm. This step is crucial to ensure the accuracy of morphology tracking over time.

To perform manual corrections, experts must observe and compare the shapes of clusters across sequential images. If the shape in a subsequent image significantly deviates from its predecessor, it may indicate that it belongs to a different group of shapes or requires the creation of a new group altogether. Conversely, shape congruence between images confirms the accuracy of the existing cluster relationships, and no changes are necessary.

As depicted in Fig. 5, in Cases 1 and 2, the relationship for Cluster 1 remains consistent over time, demonstrating evolutionary congruence. However, notable changes are observed in other clusters. For example, in Case 1, Clusters 2 and 3 transition from a ring-like shape to a blotch, while Cluster 4 evolves from a blotch to small particles.

**Figure 5.**
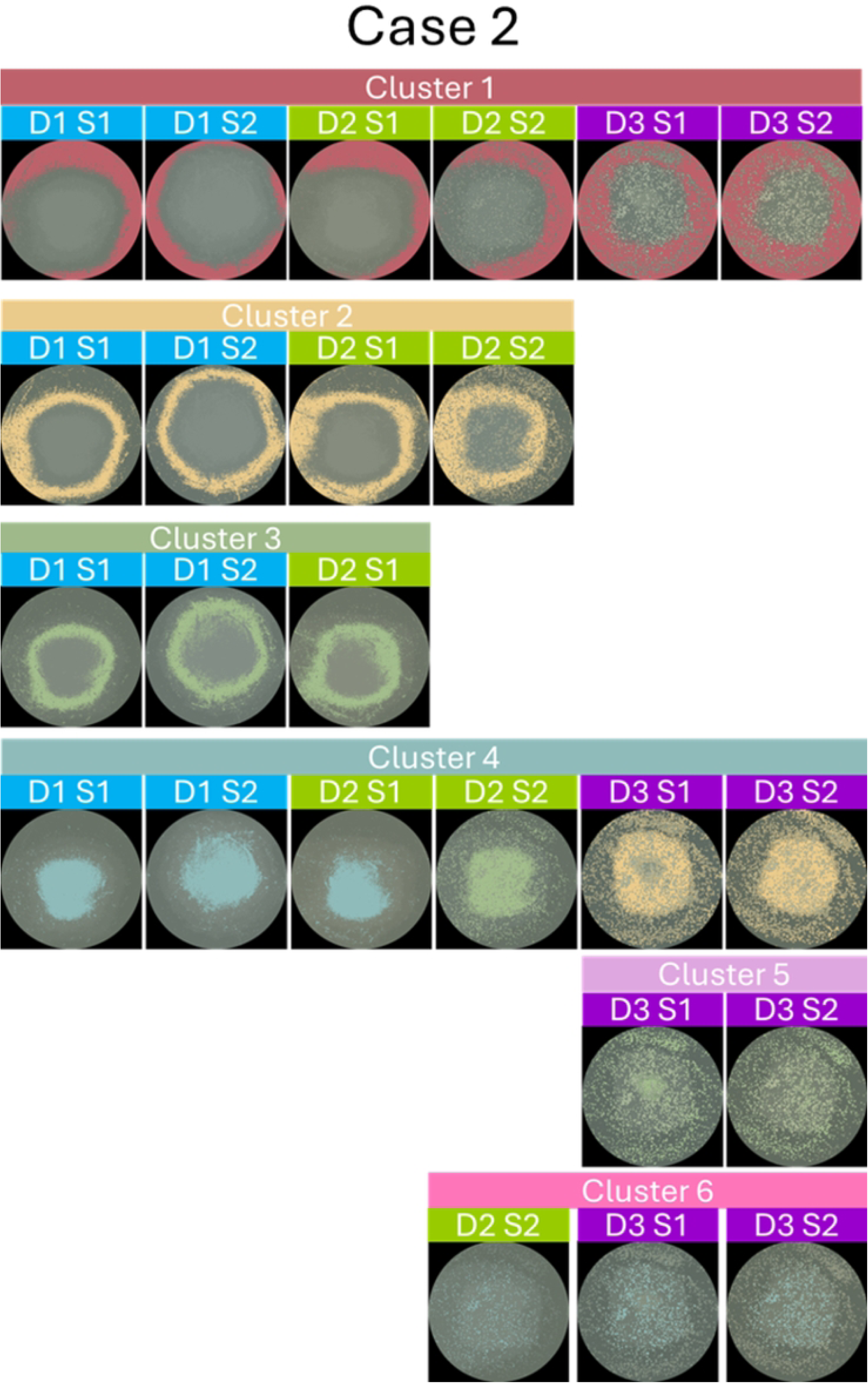
Manual corrections. Discontinuities within the same clusters are visually plausible and are indicated with red curved lines and an “x” symbol signifying incorrect correspondence along the sequence. A second correction is illustrated with green directed lines, meaning cluster correspondence.

These discontinuities are visually plausible and are indicated in the figure with red curved lines and an “x” symbol signifying incorrect correspondence along the sequence. A second correction is illustrated with green directed lines. In Case 1, these lines indicate a continuity that must be followed for a new set of clusters, starting with Cluster 4, continuing with Cluster 3, and finishing with Cluster 2. Similar observations are made for Case 2, suggesting coherence in the development of clusters for the two presented cases of *P. koreensis*. Notably, the case of *E. coli* (Case 3) did not present any discontinuities, thus no manual correction was needed to determine the final number of clusters. The consistency in shapes across images within this case implies that the original automatic clustering was accurate, and the relationships among clusters were maintained throughout.

Semi-automatic clustering requires expert observation and consequent adjustments, such as reassigning shapes to different clusters or establishing new ones. Additionally, if new groups emerge, they must be matched accordingly. The resulting final clusters, after modifications and additions, lead to the refined cluster arrangements shown in Figure 6 and Figure 7 for Cases 1 and 2, respectively. These arrangements are discussed in detail in the discussion section.

**Figure 6.**
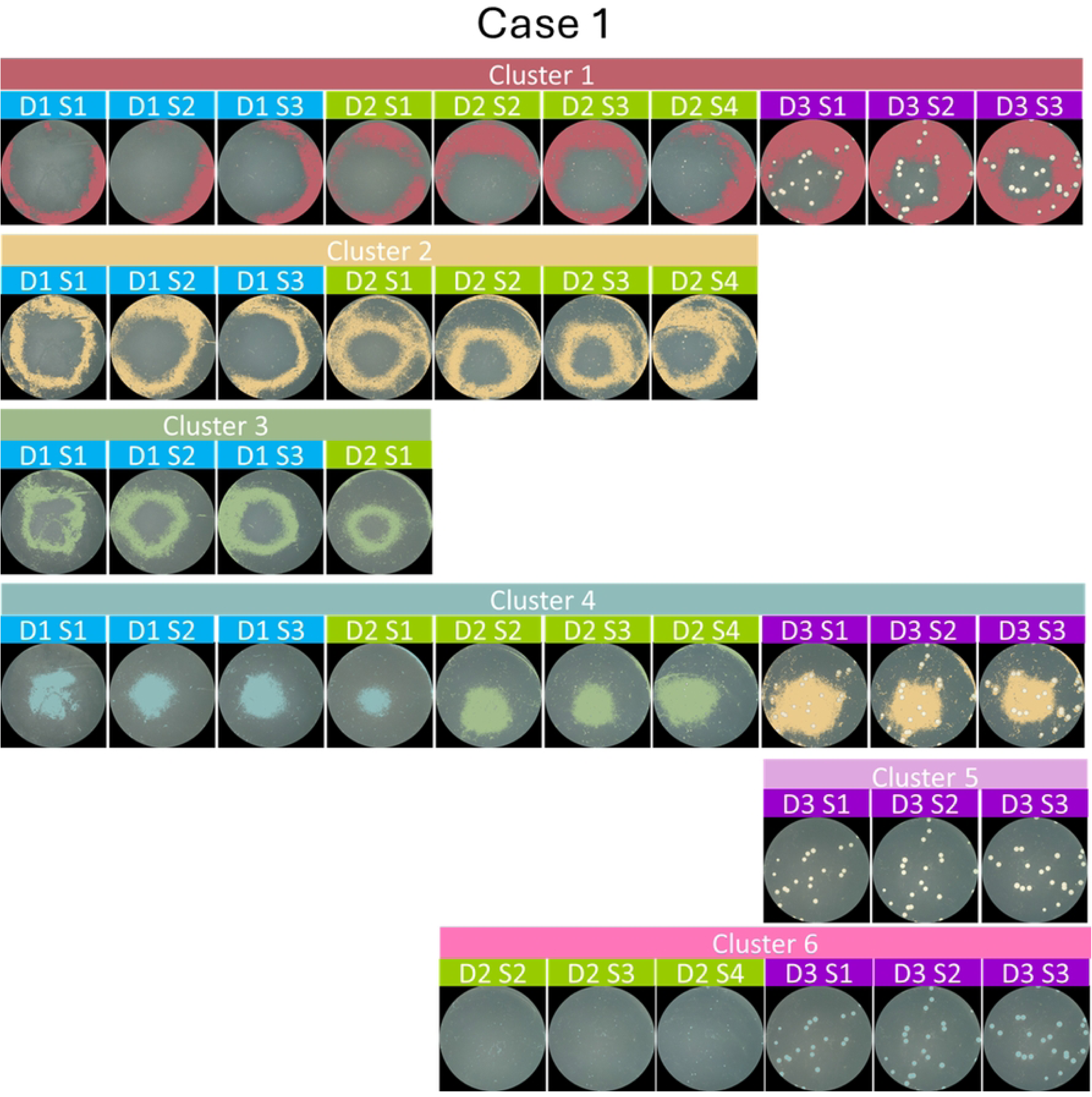
Case 1: modifications implemented with new cluster arrangement. This is the final segmentation result for Case 1. Note how Clusters 5 and 6 are added after manual correction.

**Figure 7.**
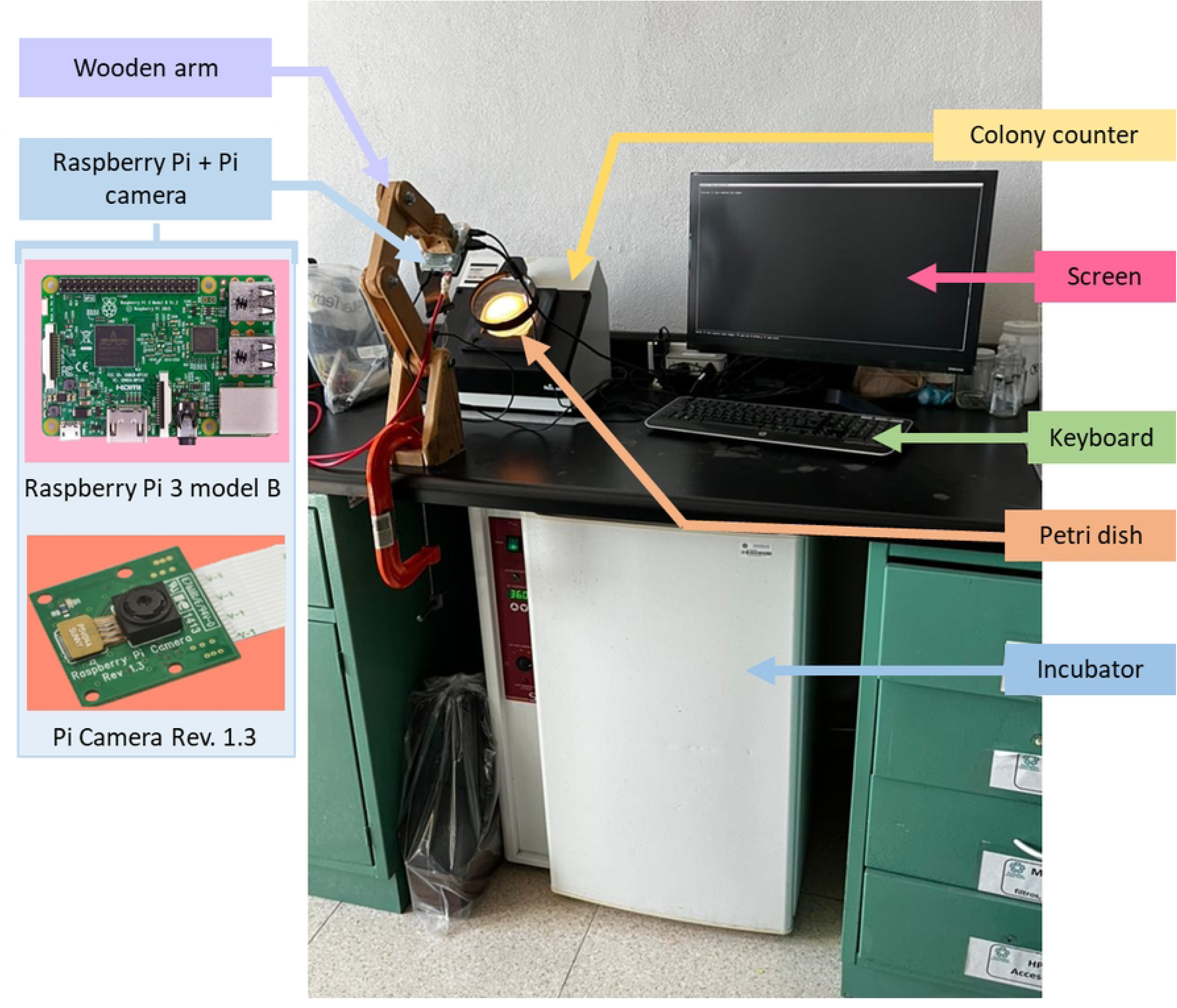
Case 2: modifications implemented with new cluster arrangement. This is the final segmentation result for Case 2. Note how, similar to Case 1, Clusters 5 and 6 are added after manual correction.

### Obtaining occupancy percentage

Following manual adjustments, the clusters are finally quantified in Python, incorporating the necessary elements to track occupancy accurately. This final stage culminates in the creation of a graph that effectively illustrates the evolution and distribution of the different morphological groups within the cultures over time.

To enhance the understanding of the experimental results, line graphs were plotted for each cluster within each case, as shown in Figures 8, 9, and 10. This is particularly effective for detailed analysis, allowing for a clear comparison of cluster dynamics over time against the standard bacterial growth curve displayed in Figure 11. Line graphs facilitate a comprehensive observation of the temporal evolution of each bacterial group within the samples, graphically representing changes in cluster size, enabling easy identification of growth trends or declines across the experimental duration. The visualization of these dynamics allows the expert to discern patterns and potentially correlate specific environmental or experimental conditions with changes in bacterial behavior. Our method not only underscores the dynamic nature of bacterial colonies but also provides a valuable tool for interpreting complex biological data in a more accessible and informative manner.

**Figure 8.**
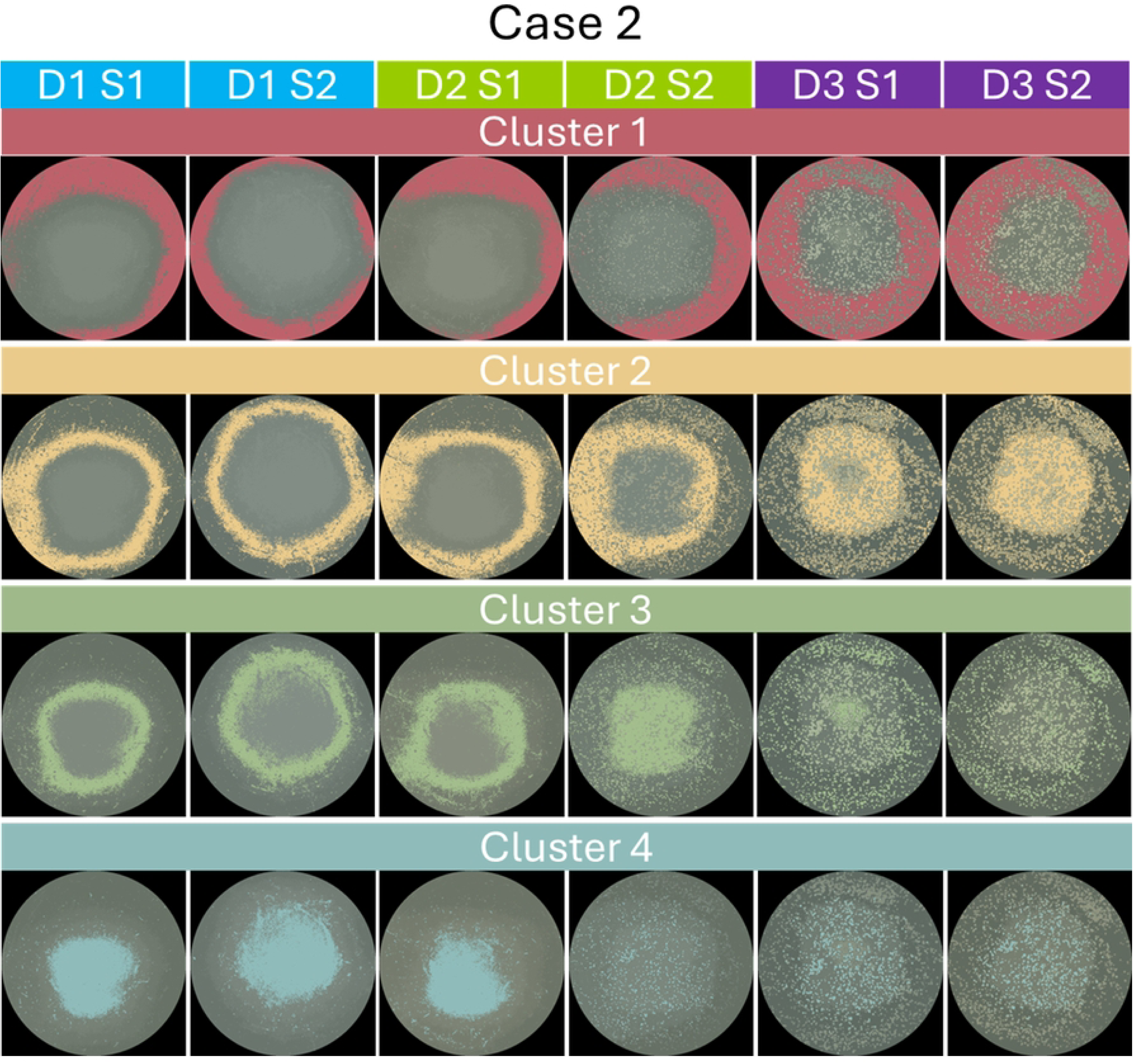
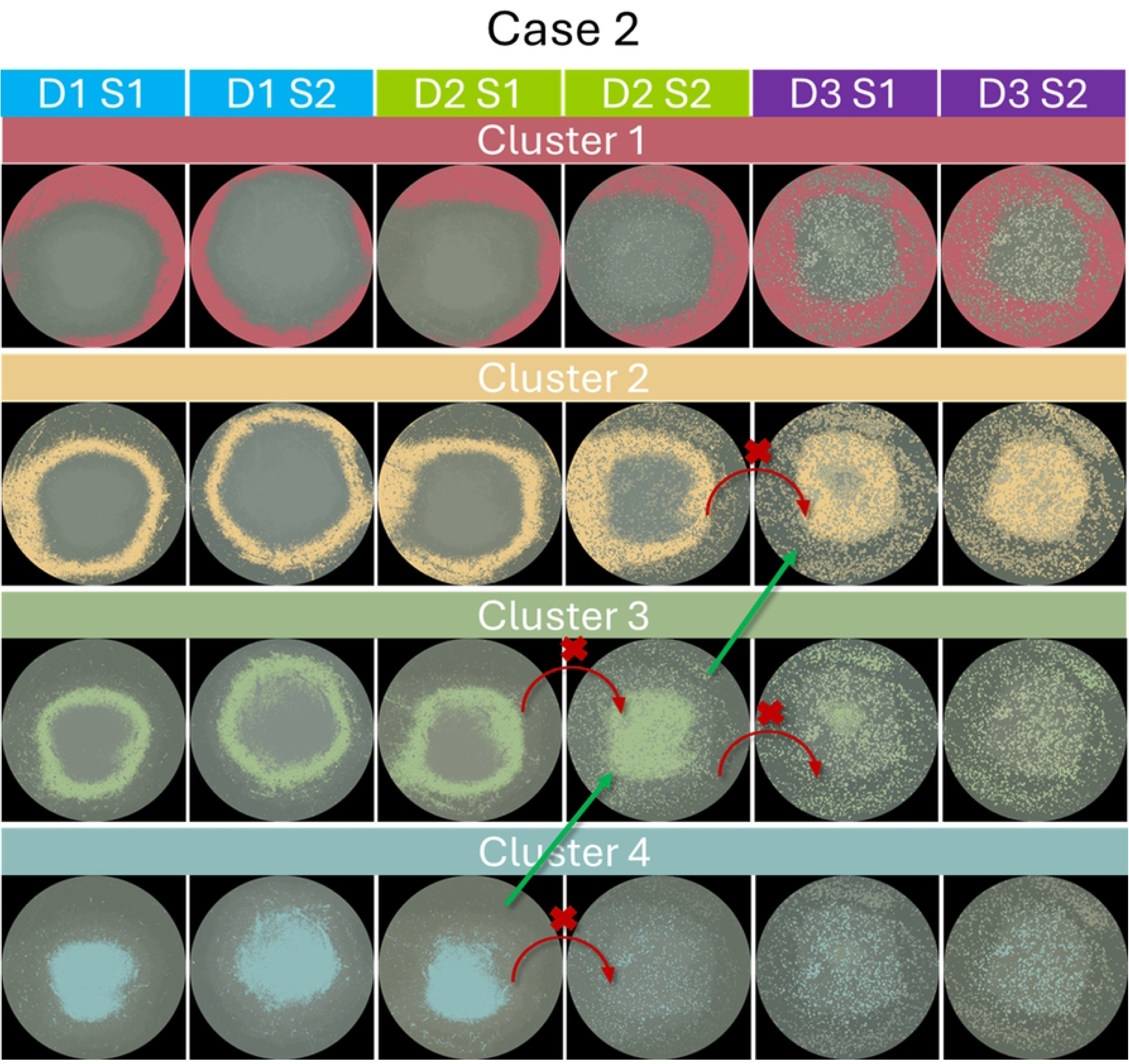
Total Occupation Percentage for Each Cluster: 1st Case. The graphs show the evolution per cluster of the ten obtained images. Note the appearance of cluster 6 after the fourth image and cluster 5 after the seventh image. Left: original data, right: smoothed data.

**Figure 9.**
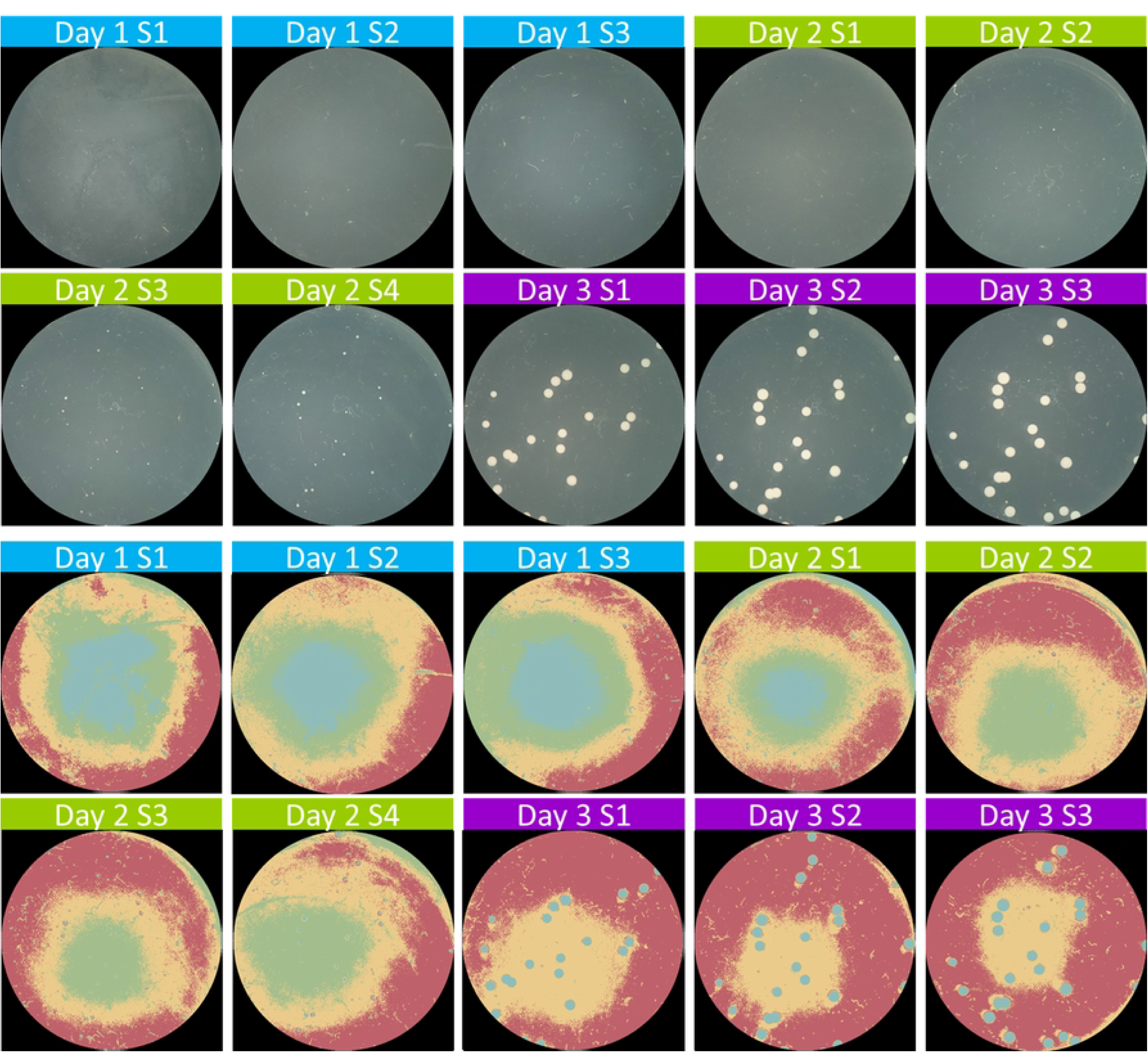
Total Occupation Percentage for Each Cluster: 2nd Case. The graph shows the evolution per cluster of the six obtained images. For this case, the appearance of cluster 6 after the third image and cluster 5 after the fourth image. Left: original data, right: smoothed data.

**Figure 10.**
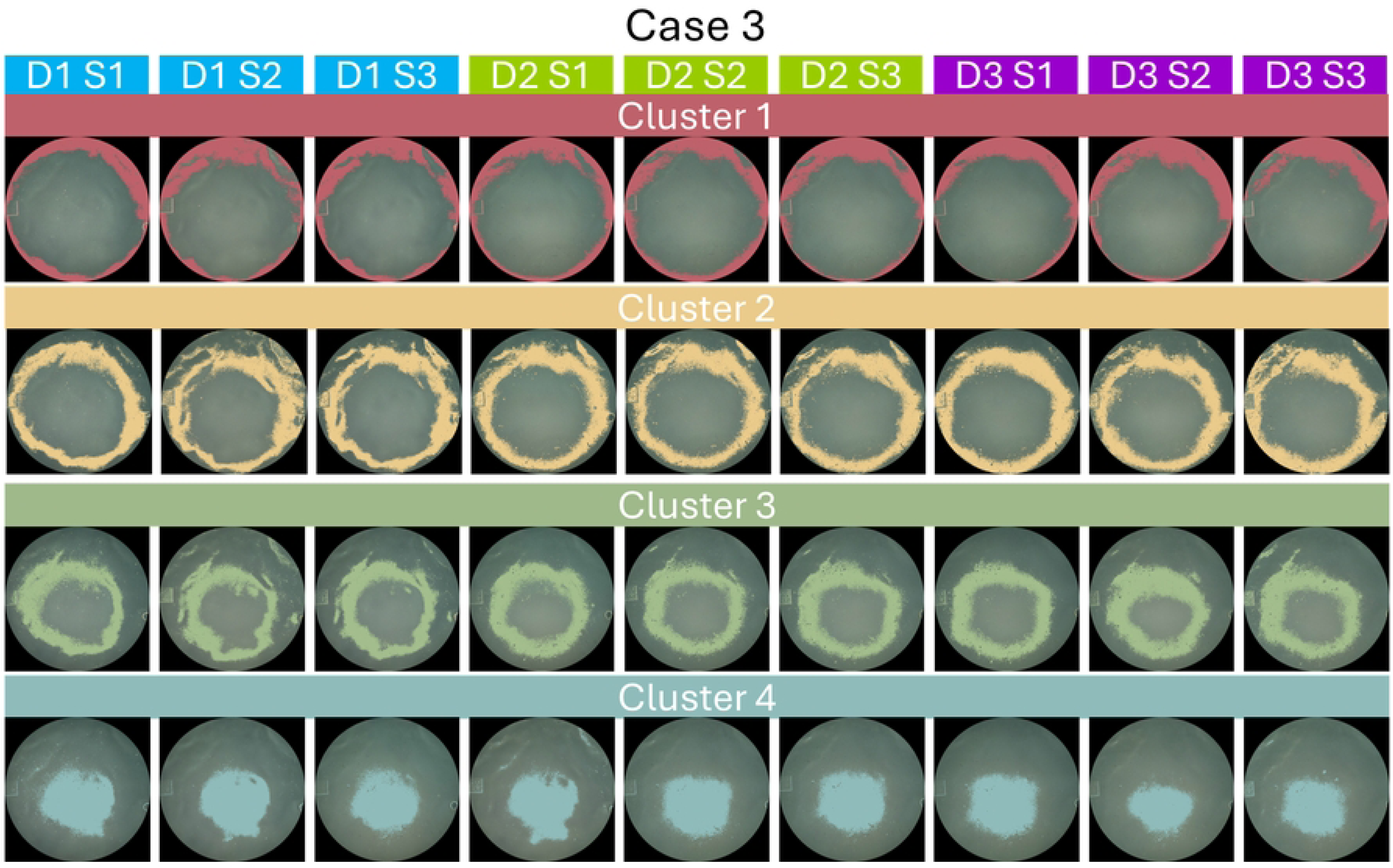
Total Occupation Percentage for Each Cluster: 3rd Case. The graph shows the evolution of each original cluster of the nine obtained images. Left: original data, right: smoothed data.

**Figure 11.**
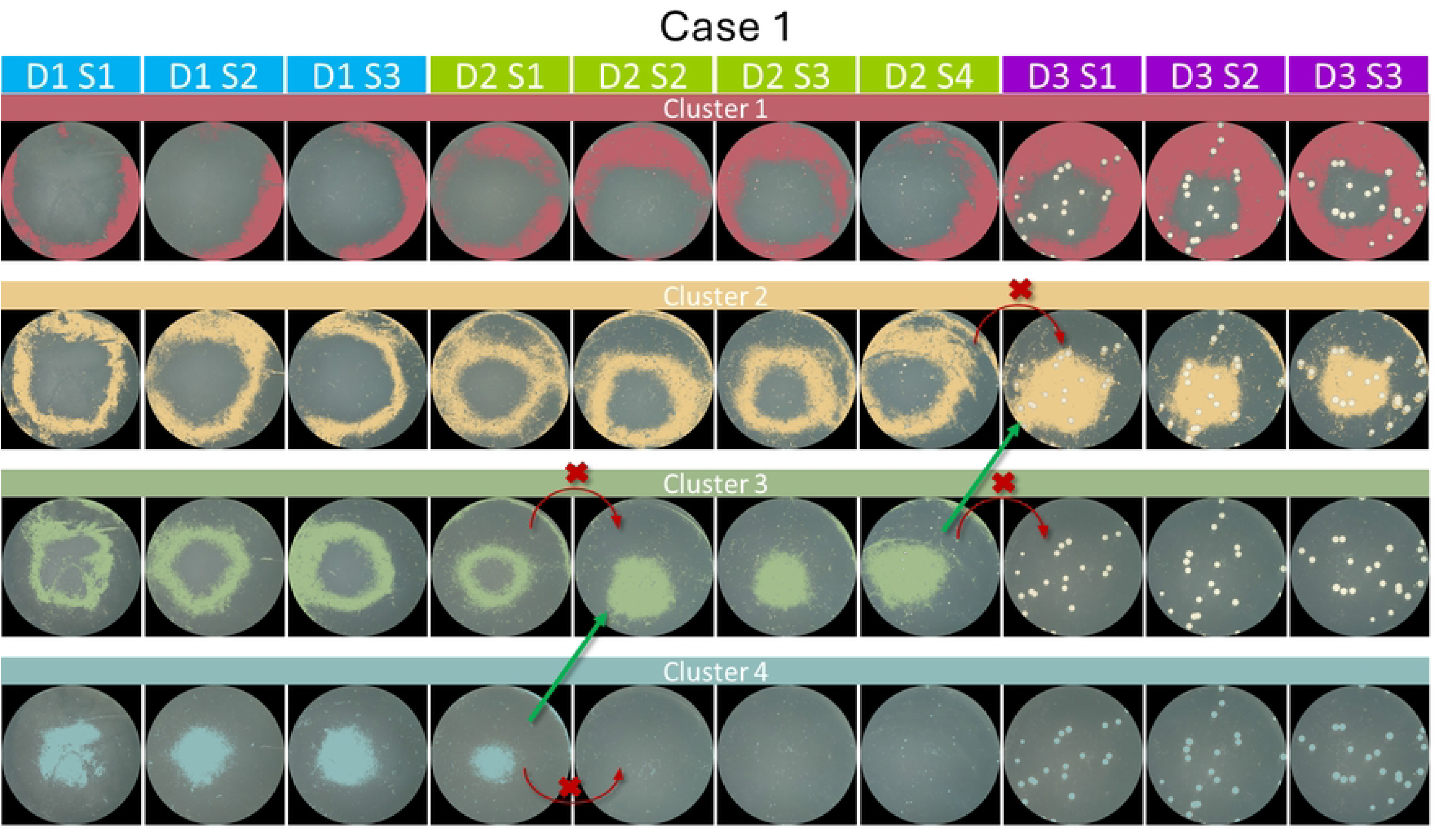
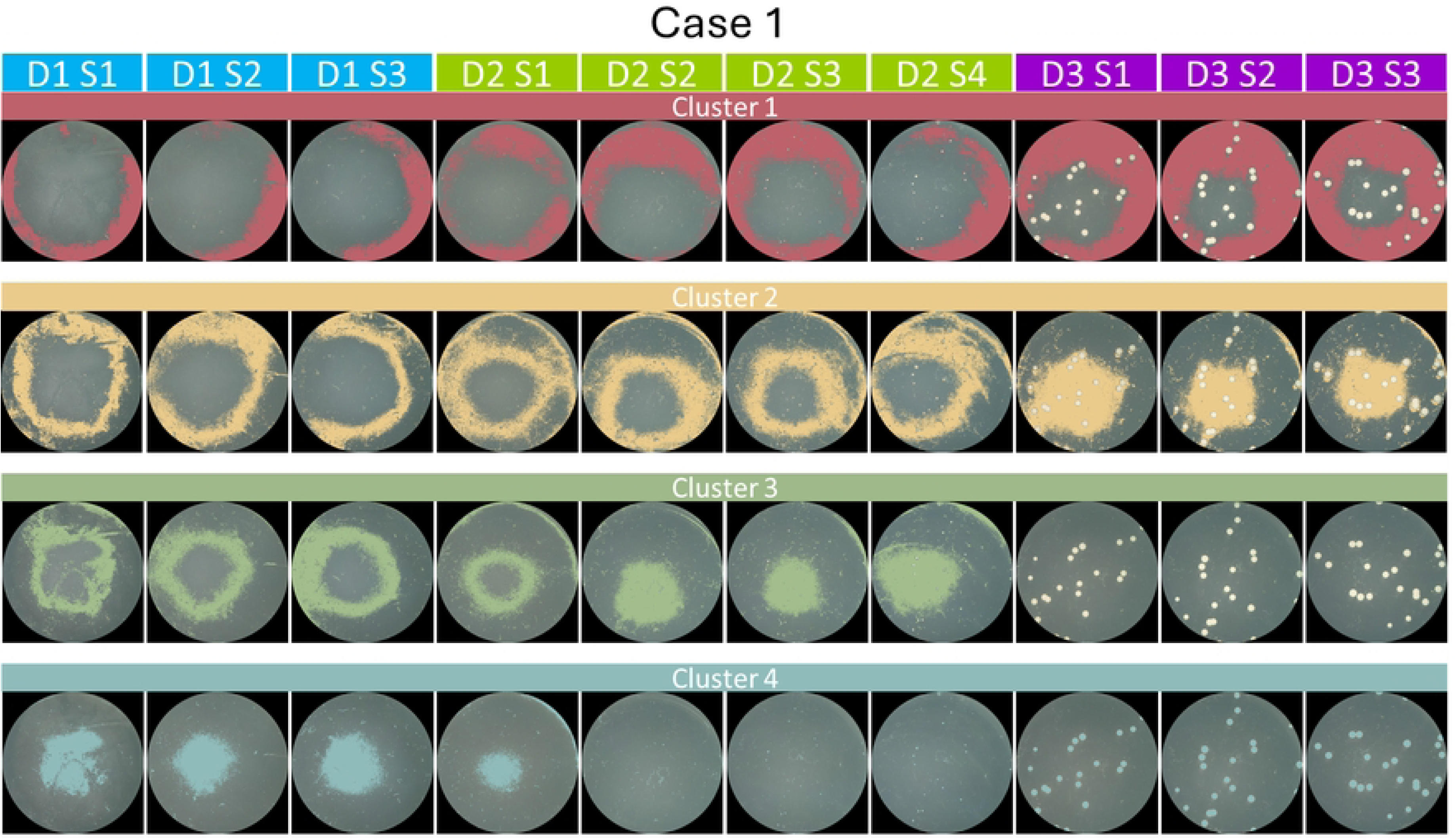
Bacterial growth curve with the four typical growth phases: Lag, logarithmic, stationary and decline phases. Based on [34].

## Discussion

In this section, we present detailed observations of each cluster from the provided cases. It is important to note that cluster 0 in each case has been consistently identified as representing the background in all images. Consequently, this cluster has been excluded from the total cluster count and subsequent analysis, as it falls outside the main area of interest —the culture— and does not contribute relevant information to the study’s objectives.

For the remaining clusters, which represent various aspects of the bacterial cultures, we will examine their evolution and significance within the context of the experiment. Each cluster’s behavior, including changes in size, shape, and color, provides insights into the dynamics of bacterial growth and interaction within the culture environment.

### Case 1

The analysis of Case 1 reveals distinct behaviors and transitions within each cluster, highlighting their roles and dynamics in the bacterial culture of *Pseudomonas koreensis*:

- **Cluster 1:** This cluster is essential as it outlines the peripheral evolution of the bacterial culture. Notably, it demonstrated a classic bacterial growth curve with phases of lag, exponential growth, and a stationary period. A significant observation from the growth curve is the sharp decline in the sixth image due to the expansion of Cluster 2. However, Cluster 1 regained prominence in the seventh image as it absorbed Cluster 2, resulting in a merged and thicker area that persisted through the remaining series of images.
- **Cluster 2:** Notably affected by manual corrections, this cluster faced challenges in consistent detection across the series. The manual edits were necessary due to the non-uniform capture of the areas by the initial automatic detection, highlighting some of the limitations of the semi-automatic method in handling dynamic changes within the culture.
- **Cluster 3:** Initially the smallest cluster, it underwent dramatic transformations starting from the third image, evolving into a distinct spot at the center of the sample. This transformation marked the genesis of a new cluster, indicating significant morphological changes within the culture.
- **Cluster 4:** Undergoing the most substantial modifications, this cluster achieved continuity with Clusters 2 and 3 across different stages of the experiment. These changes underscore the dynamic interplay between different bacterial groups within the culture.
- **Cluster 5:** As the newest addition and the smallest cluster, it represents a crucial phase in the bacterial lifecycle where punctiform bacteria begin to secrete extracellular polymers. This observation provides a novel approach to quantify and monitor such activities, which are typically assessed through labor-intensive and costly manual methods involving polymer extraction and weight measurement.
- **Cluster 6:** Emerging in the fourth image, this cluster highlights the appearance and growth of punctiform bacteria, which are often overlooked in manual counts until they manifest prominently. The detection of such bacteria early in their development is a significant advantage of the imaging and clustering techniques used in this study.

### Case 2

In Case 2, the clusters display various patterns of bacterial growth and interactions, providing further insight into the complex dynamics of microbial development for *Pseudomonas koreensis*:

- **Cluster 1:** This cluster closely mirrors a typical bacterial growth curve. It begins with moderate occupancy, progresses through exponential growth, and eventually reaches a plateau where no further significant growth occurs, marking the stationary phase. This suggests that the bacteria in this cluster have reached their environmental or nutrient limits.
- **Cluster 2:** Initially, Cluster 2 follows a typical growth curve pattern with a lag phase followed by exponential growth. However, from the fourth image onward, its space begins to merge with that of Cluster 1, illustrating the fluid nature of bacterial boundaries in a growing culture.
- **Cluster 3:** Similar to the first case, Cluster 3 is one of the smaller original clusters. From the third image, its area starts merging with that of Cluster 2 and later with Cluster 1, indicating significant overlap and interaction among these clusters.
- **Cluster 4:** This cluster exhibits a clear bacterial growth trajectory, including identifiable lag, exponential growth, and early stationary phases. Its progression most accurately represents the average growth behavior of the entire sample, providing a standard for comparison with other clusters.
- **Cluster 5:** Due to the limited number of images and the timing of the experiment, this cluster provides only a brief glimpse into the initial stages of extracellular polymer release by punctiform bacteria. Extended sampling might have offered more detailed insights into this significant biological process.
- **Cluster 6:** As in the previous case, this cluster highlights the growth of bacteria with defined shapes, emerging from the third image onward. Previously part of Cluster 4, this cluster demonstrates how distinct bacterial forms can evolve and become more prominent over time.

### Case 3

Case 3 presents unique challenges and phenomena among the clusters, showcasing the nuanced interactions and morphological changes that occur during the bacterial culture development of *Escherichia coli* :

- **Cluster 1:** This cluster exhibits significant variations in shape across the sequence of images, likely resulting from manual edits during the segmentation process. These inconsistencies suggest difficulties in maintaining stable cluster boundaries, impacting the reliability of morphological analysis for this cluster.
- **Cluster 2:** Most closely aligning with the overall bacterial growth curve of the sample, Cluster 2 shows a somewhat inconsistent lag phase in the earliest images (0-3). However, from image 3 onwards, a clear growth phase is observed, during which this cluster maintains a consistent area and achieves a high percentage of occupancy, indicating robust bacterial development.
- **Cluster 3:** The evolution of this cluster is marked by variability, starting with moderate values, then experiencing a decrease, followed by growth. Its position, in the middle of Clusters 2 and 4, results in fluctuating occupancy percentages as it is encroached upon by both neighboring clusters, possibly underscoring the competitive aspect of spatial occupancy in bacterial cultures.
- **Cluster 4:** This cluster undergoes the most substantial reduction in space over time, indicating it faces the highest level of competition or unfavorable conditions within the culture. Starting from the second image, its area progressively diminishes, ultimately occupying only 15.36% of the available space by the end of the sequence. This trend highlights the challenges some bacterial groups may encounter as they compete for resources or adapt to environmental pressures within the culture.

## Conclusion

The tracking experiments conducted in this study yielded substantive insights, demonstrating the effectiveness of the semi-automatic procedure in mirroring the typical bacterial growth curve phases—lag, exponential, and stationary—across various clusters in the three cases analyzed. This alignment not only validates the semi-automatic method but also enhances the understanding of bacterial dynamics within cultures.

The application of automatic clustering facilitated initial perceptual and manual corrections, especially highlighted in the two cases that required adjustments. These cases exhibited similar behavioral patterns and underwent comparable reorganization, showcasing the utility of automated clustering in simplifying the correction process.

This approach provided a strong foundational layer for subsequent manual interventions, proving crucial for accurate morphological analysis.

Additionally, it was observed that the consistency of the culture medium significantly affects both the bacterial growth and the segmentation accuracy. The case that required no corrections was associated with a more gelatinous agar medium, which likely contributed to a more uniform segmentation output due to less variation in bacterial morphologies and more homogeneous image characteristics. This finding highlights the influence of physical media properties on the effectiveness of image-based analytical techniques.

The use of the k-means algorithm for segmenting images has proven instrumental in revealing detailed information about bacterial morphology and concentration. This technique effectively highlighted intricate patterns and details that could be easily overlooked, even by skilled observers. By discerning subtle variations in tone and identifying mixed morphologies, a deeper and more nuanced understanding of the bacterial samples was achieved.

Furthermore, employing occupancy percentage as a metric for analyzing bacterial samples offers significant advantages over traditional Colony Forming Unit (CFU) counts. This metric considers the full morphology of the colonies, thus providing a more comprehensive evaluation that accommodates the diverse shapes and sizes of bacterial growth. This approach not only captures the quantitative aspects of bacterial cultures, but also reflects the qualitative diversity within the microbial community, making it a superior method for detailed scientific analysis in microbiology.

## Acknowledgments

The first author of this article acknowledges CONAHCYT for the support provided within its National Scholarship Program (grant number 800569). She also thanks the Robotics and Advanced Manufacturing Group for the training provided, and the Department of Sustainability of Natural Resources and Energy for the collaboration and assistance. This research has been funded by Project CONAHCYT 845101, Accelerated Discovery of Antibiofouling Materials.

## Author Contributions

Conceptualization: Diana A. Alvarado-Ruiz Mario Castelán, Keny Ordaz-Hernández, Lourdes Díaz-Jiménez.

Data Curation: Grecia L. Lara-Cadena, Roberto González-López.

Formal Analysis: Diana A. Alvarado-Ruiz, Keny Ordaz-Hernández

Funding Acquisition: Gregorio Vargas-Gutiérrez

Investigation: Diana A. Alvarado-Ruiz

Methodology: Diana A. Alvarado-Ruiz, Keny Ordaz-Hernández

Project Administration: Mario Castelán, Keny Ordaz-Hernández, Lourdes Díaz-Jiménez

Resources: Keny Ordaz-Hernández, Lourdes Díaz-Jiménez Software: Diana A. Alvarado-Ruiz, Keny Ordaz-Hernández

Supervision: Mario Castelán, Keny Ordaz-Hernández, Lourdes Díaz-Jiménez Validation: Diana A. Alvarado-Ruiz, Mario Castelán, Keny Ordaz-Hernández, Lourdes Díaz-Jiménez

Visualization: Diana A. Alvarado-Ruiz

Writing – Original Draft Preparation: Diana A. Alvarado-Ruiz

Writing – Review & Editing: Diana A. Alvarado-Ruiz, Mario Castelán, Keny Ordaz-Hernández, Lourdes Díaz-Jiménez

## Supporting Information

### Bacterial Growth Phases

In the context of our experiments, understanding the bacterial growth curve is fundamental. This curve, as shown in Figure 11, illustrates the progression of bacterial growth over time and is divided into four distinct phases, as outlined by Tortora [33]:

1. **Lag Phase:** This initial phase is characterized by little to no increase in the number of observable cells. It is termed the “lag” phase because it can vary significantly in duration, lasting from an hour to several days. Despite the apparent lack of growth, this phase is a period of intense metabolic activity where cells acclimate to their surroundings and prepare for active growth.
2. **Logarithmic (Log) or Exponential Growth Phase:** During this phase, cells begin to divide at a rapid pace, and their numbers increase exponentially. This phase features maximum reproductive activity, and the generation time—the time it takes for the population to double—reaches its minimum and stabilizes. On a logarithmic scale, the growth curve during this phase appears as a straight line due to the constant rate of exponential growth.
3. **Stationary Phase:** This phase represents a balance between cell growth and cell death, resulting in a stable population. The growth rate slows, and the total number of viable cells plateaus. Factors contributing to this phase include the depletion of nutrients, accumulation of waste products, and adverse changes in pH, all of which inhibit further growth.
4. **Decline or Death Phase:** In this final phase, the number of dying cells surpasses the number of new cells being produced. The decline continues until the population is significantly reduced or, in some cases, until all cells die. This phase results from prolonged nutrient depletion, continued accumulation of toxic waste products, and overall environmental deterioration unsuitable for sustaining bacterial life.

